# Dietary Patterns and Gene Expression Profiles in Gastric Adenocarcinoma Patients from the Northern Region of Brazil

**DOI:** 10.1101/2025.09.17.676793

**Authors:** Lidiany Corrêa Serrão, Valéria Cristiane Santos da Silva, Ana Karyssa Mendes Anaissi, Diego Pereira, Tayana Vago de Miranda, Daniel de Oliveira Mackert, Jessica Manoelli Costa da Silva, Ana Paula Freitas de Sousa, Naiza Nayla Bandeira de Sá, Williams Fernandes Barra, Fabiano Cordeiro Moreira, Paulo Pimentel de Assumpção, Samia Demachki, Marilia de Souza Araújo

## Abstract

**Background:** Gastric adenocarcinoma (GA) remains a major public health concern worldwide, with incidence influenced by demographic, socioeconomic, and lifestyle factors. Diet is a modifiable risk factor that can shape tumor biology, potentially affecting gene expression and the tumor microenvironment. Understanding how dietary patterns correlate with molecular signatures in GA may provide insights for preventive and therapeutic strategies.

**Methods:** This retrospective, cross-sectional, analytical study included 60 patients diagnosed with gastric adenocarcinoma and treated at the João de Barros Barreto University Hospital, Pará, Brazil. Structured questionnaires collected sociodemographic, lifestyle (smoking and alcohol), clinical, and dietary data. Dietary patterns were analyzed according to the Brazilian Ministry of Health guidelines. A subset of 16 patients was selected for gene expression profiling to evaluate correlations between dietary exposure and molecular alterations.

**Results:** The patient cohort exhibited a predominance of males (56.7%) and older adults (>60 years), often from socioeconomically disadvantaged backgrounds. Dietary analysis revealed insufficient intake of fruits, vegetables, and legumes, alongside high consumption of ultraprocessed foods, especially sugar-sweetened beverages. Gene expression analysis identified 48 differentially expressed genes between patients with lower (LEDRF) and higher (HEDRF) dietary risk exposure. Six genes with potential biological relevance were highlighted: MAGEA3, IL6, HCAR2, NLRP4, PGA3, and CTCFL. MEFDR patients showed overexpression of MAGEA3, IL6, and HCAR2, associated with cell proliferation, inflammation, and tumor microenvironment remodeling.

**Discussion:** The results suggest that dietary patterns may modulate gene expression and influence the tumor microenvironment in gastric adenocarcinoma. Diets rich in ultraprocessed foods and sugar may promote a pro-inflammatory and pro-oxidant microenvironment, while consumption of fruits, vegetables, and bioactive compounds appears protective, potentially regulating epigenetic, metabolic, and immunological pathways. The study underscores the importance of integrated approaches combining nutritional intervention and molecular profiling to better understand GA progression and identify potential therapeutic targets.

## Introduction

Gastric cancer (GC) is considered the fifth most common type of cancer worldwide and the fifth leading cause of cancer-related death^1^. In Brazil, it ranks fourth among the most lethal cancers in men and sixth among women, with a particularly high prevalence in the Northern region^2^. Curado et al.^3^ reported that the highest incidence of GC is found in the city of Belém, a fact that may be related to unequal access to healthcare services across regions. Rodrigues et al.^4^ further state that regional variations in GC incidence may partly reflect differences in dietary habits, host-related factors, salt intake, and, importantly, the prevalence and virulence of *Helicobacter pylori*, given the close correlation between its prevalence and GC incidence.

Although the causes are multifactorial, an individual’s metabolic status and lifestyle, particularly diet and physical activity, as well as other lifelong environmental exposures, can either protect against cancer or increase the risk of developing the disease^2,5^. The assessment of dietary patterns has gained prominence for providing a more comprehensive representation of diet and its effects, as opposed to the evaluation of isolated nutrients. This approach considers the synergistic and antagonistic effects of nutrient combinations, offering more meaningful insights into how diet influences the risk of diseases such as cancer^6,7,8,9^.

At the interface between diet and carcinogenesis, oxidative stress emerges as a critical link. This process can damage DNA molecules, alter cellular signaling pathways, and modulate the progression of several cancer types, including breast, lung, liver, colon, prostate, ovarian, and central nervous system cancers^10,11,12,13,14^. Although it is not possible to establish a direct causal relationship between current dietary intake and the development of gastric cancer, given the multifactorial nature and long latency period of the disease, analyzing dietary patterns in diagnosed patients can provide relevant contributions to public health and to the field of nutritional epidemiology. Such analyses may enable the identification of dietary markers of risk and protection, allowing the delineation of dietary profiles associated with the disease^8^.

In the context of gastric cancer, the investigation of gene expression profiles in tumor samples can help generate relevant hypotheses, including correlations with environmental factors such as dietary habits, and further reinforce the need for personalized approaches in the management and follow-up of patients.

## Materials and Methods

### Ethical aspects and inclusion criteria

Patients diagnosed with gastric cancer, specifically adenocarcinoma, aged over 20 years, of both sexes, and undergoing treatment at the João de Barros Barreto University Hospital, regardless of treatment modality, were included in the study. The research protocol involving dietary intake assessment was approved by the Research Ethics Committee of the João de Barros Barreto University Hospital (approval no. 2.844.459) and by the Oncology Research Center (approval no. 3.460.414), in compliance with Resolution 466/2012 of the National Health Council (CNS, 2012), and data collection was initiated only after approval was granted by the respective committees. For the second part of the study, 16 tumor tissue samples obtained from patients with gastric adenocarcinoma who underwent surgical resection were analyzed, under approval of the Ethics and Research Committee of the João de Barros Barreto University Hospital (approval no. 47580121.9.0000.5634), in accordance with the principles of the Declaration of Helsinki. Participant recruitment and sample collection took place between July 2, 2022, and July 6, 2023, and all individuals were provided with detailed information about the study objectives, potential benefits, risks, and possible harms, after which written informed consent was voluntarily obtained prior to inclusion.

Method for assessing dietary intake and dietary markers selected for this study General dietary intake was assessed using a Quantitative Food Frequency Questionnaire, previously validated in Brazil, comprising over 120 food items, including regional foods from Pará. Foods were categorized according to the classification in the Food Guide for the Brazilian Population into fresh or minimally processed foods, processed foods, ultra-processed foods, and processed culinary ingredients (Brazil, 2014). The questionnaire was designed to facilitate data collection, with predefined portion sizes based on household measures or food units. Food consumption frequency was recorded as the number of times each item was consumed per day, week, month, or year, or classified as “never/rarely,” and portion sizes were categorized as small (S), medium (M), or large (L). Quantitative analysis of food intake for each group was calculated using the following formula:

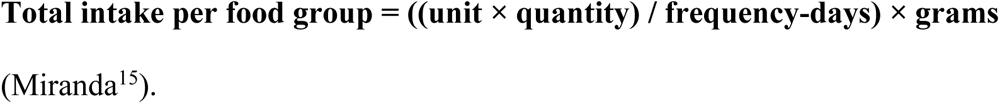

For the dietary intake analysis of all patients in this study (n = 60), only the dietary markers specified in the updated form adopted by the Primary Health Care Department of the Ministry of Health^16^ were extracted from the main project database. This form, designed for children over 2 years, adolescents, adults, elderly, and pregnant women, is based on the Food Guide for the Brazilian Population (2014) and defines healthy markers as the consumption of fruits, vegetables, and beans, and unhealthy markers as the intake of processed meats, sugary drinks, instant noodles, savory biscuits, and sweets, for characterizing the dietary profile of the sample. In the second stage, dietary data from 16 patients were selected, and based on questionnaire responses, participants were stratified into two groups: Group 0 and Group 1, representing lower and higher exposure to dietary risk factors, respectively. Foods were classified as protective (beans, fruits, vegetables) or as associated with increased risk of gastric adenocarcinoma (red meats, smoked, charcoal-grilled, processed, and ultra-processed foods), following recommendations from INCA^17^ and the Ministry of Health^16^.

### Statistical analysis of sociodemographic, lifestyle and dietary intake data

Sample characterization was performed through sociodemographic variables, categorized as: biological sex (male, female), age group (27–35, 36–43, 44–51, 52–59, 60–67, 68), educational level (illiterate, <5 years, 6–8 years, 9–12 years, high school, higher education), self-declared ethnicity (mixed-race, Black, White), place of origin (capital/interior); family history of cancer (yes/no). Lifestyle habits included tobacco and alcohol consumption, which were evaluated qualitatively (yes/no) and included only in descriptive analyses.

To assess differences in food group intake among patients in the last 24 months, the Kruskal-Wallis test, for nonparametric data, was used to compare medians of consumption frequency across different food groups, both in the total patient group and in subgroups by gender (female and male). The Dunn test was used to identify specific groups with significant differences. Statistical analyses were performed using R software version 4.2, with data of interest selected and tabulated in Excel 10.

### Statistical analysis of dietary score data

For correlation with gene expression analysis, dietary data were processed through modeling and risk calculation, based on epidemiological studies and probabilistic models, which combine known risk factors with the probability of developing a specific type of cancer^18^. Weights were assigned to each patient’s response, allowing classification into risk categories and calculation of a dietary exposure score.

From this score, patients were divided into two groups:

● Group 0 (LEDRF): dietary profile with lower exposure to dietary risk factors, based on consumption of fruits, vegetables, legumes, and beans, with scores between -7 and 0.
● Group 1 (HEDRF): dietary profile with higher exposure to dietary risk factors, based on total consumption of ultra-processed foods, soft drinks or sugary drinks, red meats, cured, smoked, charcoal-grilled meats, with scores between 0 and 7.

It is important to highlight that this classification does not refer to the future probability of disease development, as all patients were already diagnosed, but rather to a retrospective estimate based on usual dietary patterns, allowing investigation of possible correlations between dietary profile and tumor gene expression.

### Gene expression evaluation of patients

To evaluate patient gene expression, dietary intake data were integrated with gene expression data from the project *“Characterization of potential biomarkers and therapeutic targets of Gastric Adenocarcinoma in the state of Pará”*.

### Total RNA Extraction and Sequencing

Approximately 30 mg of tumor tissue from each sample was macerated for RNA extraction using TRIzol Reagent (Thermo Fisher Scientific) according to the manufacturer’s instructions. Total RNA integrity and concentration (ng/µL) were assessed with the Qubit 4 Fluorometer (Thermo Fisher Scientific), with RNA integrity number (RIN) ≥ 5 as the inclusion criterion. Library preparation was performed using the TruSeq Stranded Total RNA Library Prep Kit with Ribo-Zero Gold (Illumina, US), following the manufacturer’s protocol, with approximately 1 µg of total RNA per sample in a final volume of 11 µL. After library construction, fragment integrity of the generated DNA was evaluated using the 2200 TapeStation System (Agilent Technologies AG, Basel, Switzerland). Sequencing was conducted in paired-end mode on the NextSeq® 500 platform (Illumina®, US) with the NextSeq® 500 MID Output V2 Kit – 150 cycles, following the manufacturer’s instructions.

### Quality Control of Reads and Alignment with Human Genome

After sequencing, RNA-Seq reads were subjected to quality control. Initially, read quality was assessed using FastQC. Subsequently, adapters and low-quality reads were removed using Trimmomatic v0.39, with a quality value (QV) threshold set at >15.

Human transcript reads were characterized through alignment and quantification with Salmon v1.5.2, using coding transcripts from hg38 (www.ensembl.org) as reference index and GENCODE v42 (www.gencodegenes.org) as annotation. The reads quantified by Salmon were imported into RStudio using the Tximport v3.14.0 package.

### Differential Gene Expression Analysis

After alignment, differential gene expression profiling was performed between patients with lower exposure to dietary risk factors (LEDRF) and those with higher exposure (HEDRF).

### Functional Enrichment Analysis of Gene

Differentially expressed genes were enriched using Gene Ontology (GO) for terms related to biological processes, aiming to better understand their biological and functional relevance. The ClusterProfiler v4.8.3 package was used for this analysis.

## Results

### Patient characterization

A total of 60 patients diagnosed with gastric adenocarcinoma were evaluated, of whom 56.7% were biologically male, with the majority self-identifying as Brown/mixed-race (86.7%) and a smaller proportion as Black (8.3%). Most cases were concentrated in individuals aged between 59 and 75 years, and more than half of the participants (55%) originated from rural areas rather than the capital. A considerable proportion presented low educational levels, with 43.3% reporting fewer than eight years of schooling. Lifestyle factors included tobacco use in 53.3% of patients and regular alcohol consumption in 50%, while approximately 30% reported a family history of cancer involving different tumor types (Table 1).

**Table 1.** Demographic and lifestyle data of patients with gastric adenocarcinoma treated at HUJBB.

| Variable | Absolute frequency (n) | % |
| --- | --- | --- |
| **Biological sex** |  |  |
| Male | 34 | 56.67 |
| Female | 26 | 43.33 |
| **Age group** |  |  |
| 27–35 | 1 | 1.69 |
| 36–43 | 6 | 10.17 |
| 44–51 | 8 | 13.56 |
| 52–59 | 6 | 10.17 |
| 60–67 | 20 | 33.9 |
| 68–75 | 18 | 30.51 |
| **Educational level** |  |  |
| Illiterate | 6 | 10 |
| Less than 5 years | 16 | 26.67 |
| 6–8 years | 26 | 43.33 |
| 9–12 years | 3 | 5 |
| High school | 9 | 15 |
| **Origin** |  |  |
| Capital | 27 | 45 |
| Rural areas | 33 | 55 |
| **Family history of cancer** |  |  |
| Yes | 18 | 30 |
| No | 42 | 70 |
| **Self-reported ethnicity** |  |  |
| Brown/mixed-race | 52 | 86.67 |
| Black | 5 | 8.3 |
| White | 3 | 5 |
| **Smoking** |  |  |
| Yes | 32 | 53.33 |
| No | 28 | 46.67 |
| **Alcohol consumption** |  |  |
| Yes | 30 | 50 |
| No | 30 | 50 |

### Assessment of dietary markers in patients

Table 2 presents the profile of daily food consumption, in grams per day (g/day), among patients with gastric adenocarcinoma, based on the Ministry of Health criteria for healthy and unhealthy foods. Among healthy dietary markers, the mean consumption of beans was 101.15 g/day. Fruits had an average of 293.93 g/day, while vegetables and leafy greens totaled an average of 144.32 g/day.

With respect to unhealthy foods, the mean consumption of processed meats and sausages (PMS) was 37.91 g/day. Savory ultra-processed foods (SUPF) had a mean intake of 17.94 g/day. Sugar-sweetened beverages and foods (SSBF) showed a mean consumption of 484.6 g/day, with a high standard deviation of 575.48 g, indicating wide variability among individuals. For the calculation of total ultra-processed food intake, items from the database, such as plain biscuits and sweeteners, were included in addition to those specifically selected for this study. The overall mean consumption of ultra-processed foods was 540.2 g/day, with a standard deviation of 590.1 g/day.

**Table 2.** Profile of consumption of healthy and unhealthy dietary markers in patients with gastric adenocarcinoma.

| Dietary markers (g/day) | Mean | SD | Median |
| --- | --- | --- | --- |
| **Healthy** |  |  |  |
| Beans | 101.15 | 135.84 | 67.57 |
| Vegetables and leafy greens | 144.32 | 123.89 | 111.45 |
| Fruits | 293.93 | 541.72 | 113.97 |
| **Unhealthy** |  |  |  |
| Processed meats and sausages (PMS) | 37.91 | 52.61 | 24.7 |
| Savory ultra-processed foods (SUPF) | 17.94 | 71.84 | 5.26 |
| Sugar-sweetened beverages and foods (SSBF) | 484.6 | 575.48 | 241 |
| Total ultra-processed food intake | 540.2 | 590.1 | 330.73 |

Table 3 summarizes the comparative frequency of consumption of the food groups investigated in this study, considering the total patient population and stratification by gender. Since the data did not present a normal distribution, they were analyzed using the Kruskal-Wallis test, followed by Dunn’s test for multiple comparisons, with the consumption median adopted as the statistical parameter.

In the overall analysis, bean consumption was significantly higher compared to processed meats and sausages (PMS) and savory ultra-processed foods (SUPF), both with p < 0.05. On the other hand, bean consumption was significantly lower than that of sugar-sweetened beverages and foods (SSBF), also with p < 0.05. Vegetables showed a considerable intake, being significantly higher than PMS and SUPF (p < 0.05 for both).

In gender-based comparisons, women had slightly higher vegetable intake than men, although without statistical significance. Among men, fruit consumption was significantly higher compared to PMS and SUPF (p < 0.05), suggesting greater relative adherence to healthy dietary markers in this subgroup.

**Table 3.** Comparative summary of dietary marker consumption, processed meats and sausages (PMS), savory ultra-processed foods (SUPF), and sugar-sweetened beverages and foods (SSBF), 2025.

| Kruskal-wallis | Patients (n) | Beans | Vegetables | Fruits | PMS | SUPF | SSBF | Post-hoc (dunn) |
| --- | --- | --- | --- | --- | --- | --- | --- | --- |
| All patients | 60 | 67.57 | 111.45 | 113.97 | 24.7 | 5.26 | 241 | Beans × PMS - $p < 0.05$<br>Beans × SUPF - $p < 0.05$<br>Beans × SSBF - $p < 0.05$<br>Vegetables × PMS - $p < 0.05$ |
| Female patients | 26 | 46 | 114.62 | 67.83 | 20.47 | 6.58 | 205.99 | Vegetables × SUPF - $p < 0.05$<br>Vegetables × SSBF - $p < 0.05$ |
| Male patients | 34 | 86 | 105.31 | 133.71 | 30.23 | 5.26 | 337.2 | Vegetables × SSBF - $p < 0.05$<br>Fruits × PMS - $p < 0.05$ |

### Exploratory analysis of differential gene expression in collected samples

To investigate the influence of dietary patterns on the molecular profile of tumors, a comparative analysis of gene expression was performed in 16 patients, divided into two groups: Group 0, and Group 1. This analysis resulted in the identification of 48 differentially expressed genes, with emphasis on those overexpressed (log2FC > 1 and adjusted p-value < 0.05) and underexpressed (log2FC < –1 and adjusted p-value < 0.05) in each group (Figure 1).

**Figure 1.**
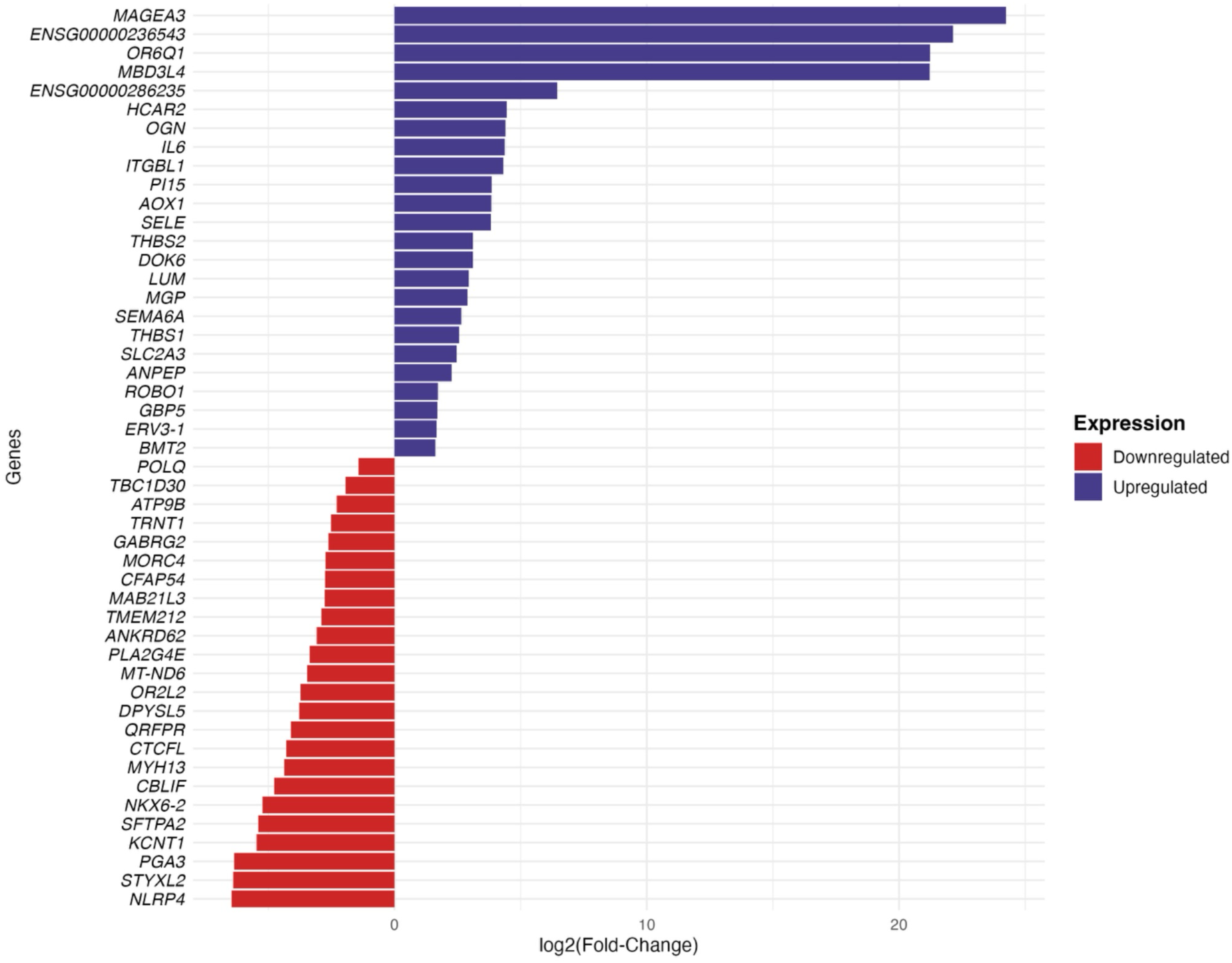
Barplot of differentially expressed genes in the comparison between LEDRF vs HEDRF groups.

To better understand the functions and influences of the differentially expressed genes between the evaluated groups, Gene Ontology (GO) analysis was performed using the ClusterProfile package, which identified metabolic and biological pathways significantly associated with the genes. Among the most representative pathways were diseases of metabolism and diseases of glycosylation, which showed the highest gene counts and statistical significance (adjusted p < 0.01). Other relevant pathways included processes related to keratin sulfate metabolism, protein glycosylation, and the metabolism of water-soluble vitamins and cofactors (Figure 2).

**Figure 2.**
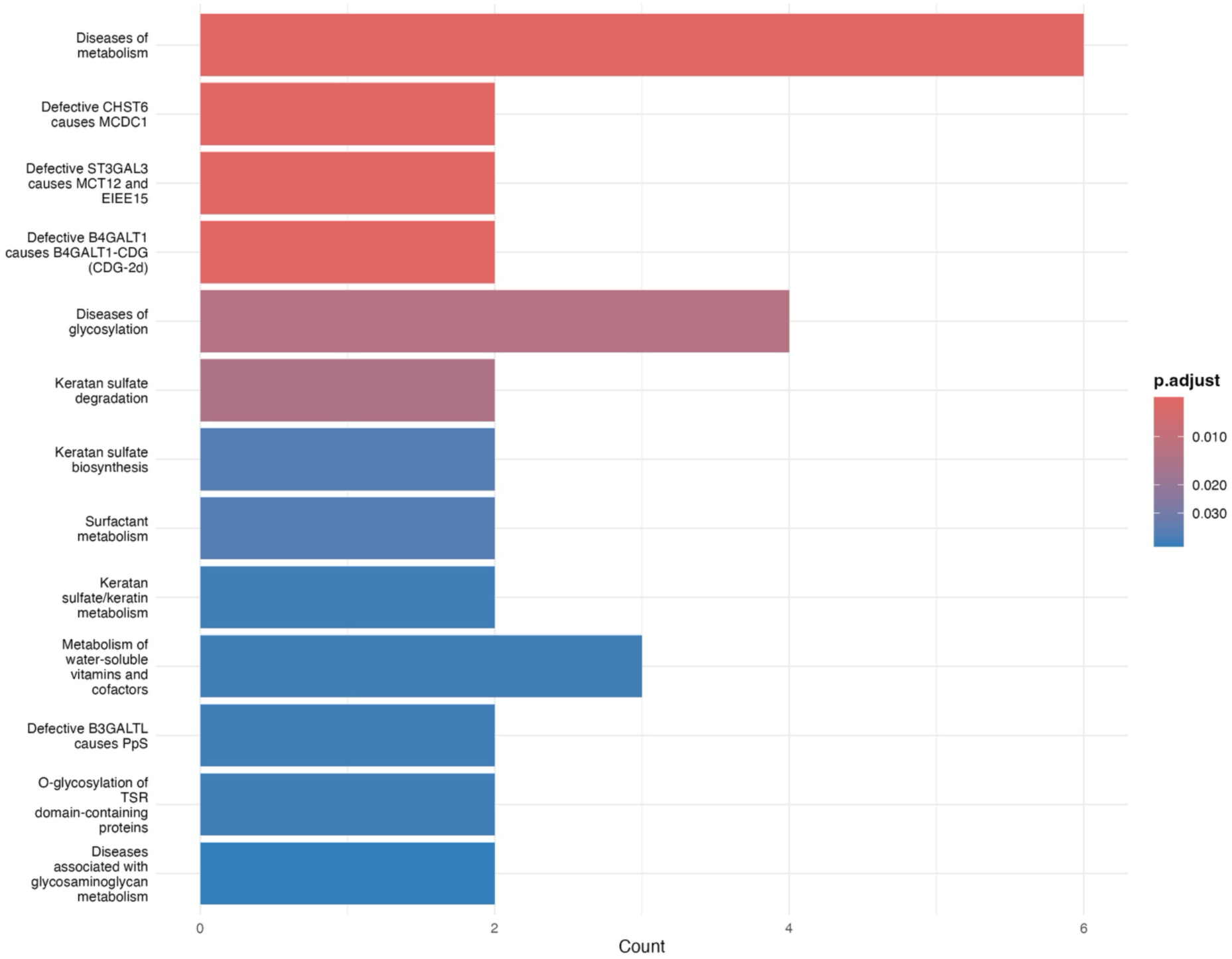
Functional enrichment analysis of differentially expressed genes.

10 genes showing differential expression were selected in Group 0: MAGEA3, OR6Q1, MBD3L4, HCAR2, OGN, IL6, ITGBL1, PI15, and 2 unidentified genes; and 10 genes in Group 1: NLRP4, STYXL2, PGA3, KCNT1, SFTPA2, NKX6-2, CBLIF, MYH13, CTCFL, QRFPR. 6 genes stand out for their biological and statistical relevance, closely related to gastric cancer and diet-associated inflammation: MAGEA3, IL6, HCAR2, NLRP4, PGA3, and CTCFL. The first 3 were significantly overexpressed in the LEDRF group, while the latter three were underexpressed in the HEDRF group.

**Figure 3.**
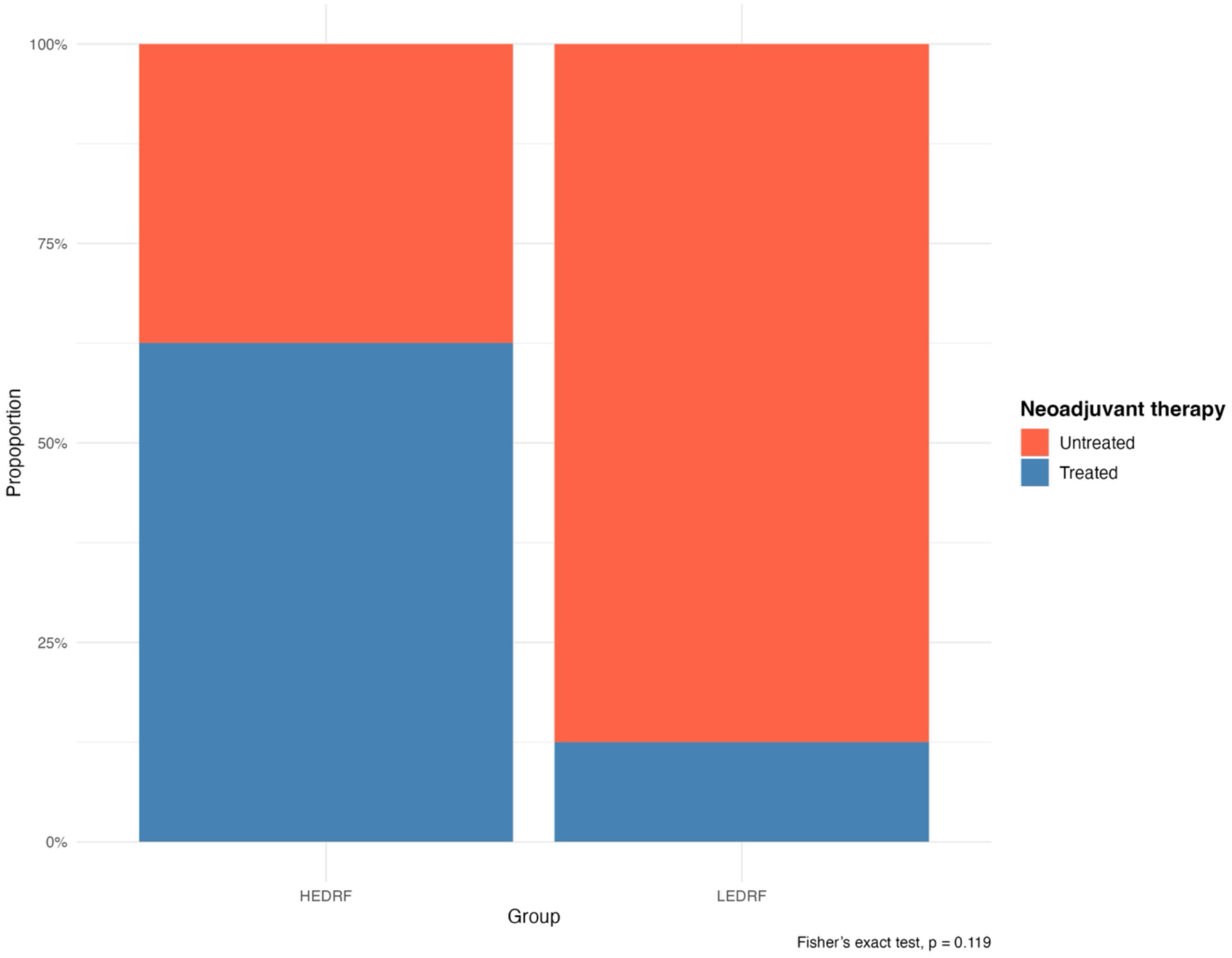
Comparison of HEDRF and LEDRF groups regarding receipt of neoadjuvant treatment.

The comparison of the proportion of patients who received neoadjuvant treatment between the HEDRF and LEDRF groups showed no statistically significant difference according to Fisher’s exact test (p = 0.119). Although not significant, there is a tendency for the HEDRF group to have a higher proportion of treated patients.

Differential gene expression analysis between patient groups classified according to dietary profile revealed distinct molecular signatures. In the LEDRF group, characterized by lower exposure to ultra-processed foods, MAGEA3, IL6, and HCAR2 were overexpressed, genes related to cell proliferation, inflammation, and tumor microenvironment remodeling. In contrast, in the HEDRF group, composed of patients with higher exposure to dietary risk factors, reduced expression levels were observed for NLRP4, PGA3, and CTCFL.

## Discussion

In this study, it was observed that patients with gastric adenocarcinoma presented an epidemiological profile that partly aligns with previously described regional and global patterns, as shown in Table 1. The predominance of 56.67% of cases occurring in male patients and the age group above 60 years, with the highest concentration of cases between 59 and 75 years, corroborates findings from other investigations that report a higher incidence of gastric adenocarcinoma in men and older individuals^1,19^.

An important finding concerns the socioeconomic profile of the participants, who were predominantly self-identified as mixed race (parda, 86.67%). A considerable proportion had a low educational level, with 43.33% reporting less than eight years of formal schooling. Moreover, more than half of the patients (55%) originated from inland municipalities, a pattern that underscores the influence of social and geographic vulnerabilities on this population. This set of characteristics suggests the influence of social determinants of health, reflecting structural inequalities and barriers to access diagnostic and treatment services, as discussed by Júnior et al.^20^ and Bray et al.^1^.

The high proportion of patients with limited education, including illiterate individuals and those with less than eight years of schooling, combined with the higher frequency of cases among residents from inland regions, reinforces the hypothesis that socioeconomic, cultural, and geographic factors may negatively impact early detection and clinical management of gastric adenocarcinoma, contributing to persistent unfavorable outcomes in more vulnerable populations.

Supporting this perspective, Jardim et al.^21^, when analyzing regional variations in cancer incidence and mortality in Brazil, showed that the incidence/mortality ratio for gastric cancer in 2018 was lower in the North and Northeast regions, suggesting poorer clinical outcomes and indicating structural inequalities in access to prevention, early diagnosis, and adequate treatment, resulting in less favorable prognoses in these areas.

The absence of a family history of cancer in 70% of evaluated patients suggests that, in this group, environmental and behavioral factors may have played a more relevant role in the onset and progression of gastric adenocarcinoma. This finding reinforces the importance of considering social determinants of health, as well as prolonged exposure to risk habits such as inadequate diet, smoking, and alcohol consumption, as central elements in the causal chain of the disease.

Regarding lifestyle, there was a high prevalence of smoking (53.3%) and alcohol consumption (50%) among the patients, although a decrease in smoking prevalence has occurred among Brazilians, especially in the North and Northeast regions^3^, both considered important habits to consider in gastric carcinogenesis^13^.

In this context, Bray et al.^1^ emphasize that the current epidemiological transition of the most prevalent cancer types, from tumors predominantly associated with infection and poverty to neoplasms related to lifestyle and diet, requires a revision of oncology control strategies, highlighting the need for public health policies to be evaluated based on new patterns.

The analysis of dietary intake patterns revealed heterogeneous habits among patients, reflecting the complexity of diet in the prevention of gastric adenocarcinoma. The markers considered healthy were fruits, vegetables, legumes, and beans, while unhealthy markers included processed and cured meats (PCM), sugar-sweetened beverages and sweets (SSB), and representatives of savory ultra-processed foods (SUPF), contained within the ‘prudent’ and ‘Western’ dietary patterns, respectively, as reported in studies by Mehta et al.^22^ and Pu et al.^8^.

Overall, the intake of healthy food markers exhibited substantial variability, as shown in Table 2, indicating highly heterogeneous dietary habits among patients. The average consumption of vegetables and greens (144.32 g/day) and fruits (293.33 g/day) stood out compared to bean consumption (101.15 g/day). For this study, the intake of potato, cassava, yam, and similar tubers was not considered. Although the average fruit consumption was significant (293.93 g), the high standard deviation (541.72 g/day) indicates large variability among patients. It is important to note that the average bean intake is considerable, showing that this staple food is part of the daily diet of many patients.

The results indicate a trend toward lower consumption of fresh foods compared to ultra-processed foods, a pattern previously identified in other studies^19,23^. This finding is particularly concerning in light of FAO/WHO recommendations, which suggest a minimum intake of 400 g/day of fruits and vegetables (excluding tubers) as an effective strategy for the prevention of various chronic diseases, including cancer^24^. Considering the high variability observed among the patients in this study, it is likely that a significant portion of the sample did not reach this recommended intake.

This variability represents a critical point, especially since studies such as Ferro et al.^7^ highlight a dose-dependent protective effect, with increasing benefits as daily portions of fruits and vegetables increase, ideally between six and ten servings per day. Moreover, recent cohort data suggest that fruits and vegetables of specific colors, such as white, red, and purple, may offer differentiated protection against gastrointestinal neoplasms^25,26^, emphasizing the importance not only of quantity but also of diversity and quality of these foods in the diet.

Supporting this evidence, a meta-analysis by Naemi Kermanshahi et al.^27^ showed that regular consumption of fruits and vegetables is associated with a reduced risk of gastric adenocarcinoma. The study found a stronger protective association for fruits than for vegetables, highlighting that every 100 g/day increase in fruit intake reduced the risk of developing the neoplasm by 5%.

These findings underscore the importance of adequate and regular intake of fresh foods as an effective preventive strategy against gastric adenocarcinoma. Conversely, the consumption of unhealthy foods observed warrants special attention (Table 2). In this study, the mean daily intake of processed and cured meats (PCM) was 37.91g, with a standard deviation of 52.61g, indicating relatively small variation among participants. This is particularly relevant when compared to the meta-analysis by Kim et al.^26^, which reported that an increase of just 50 g/day in processed meat intake is associated with a 72% increase in gastric cancer risk. Consumption of savory ultra-processed foods (SUPF) was lower, averaging 17.94 g/day.

However, the most alarming data concerns sugar-sweetened beverages and foods (SSB), with a mean intake of 484.6 g/day and a median of 241 g/day, showing high dispersion and suggesting that a significant portion of patients consumed these products in high amounts. These patterns reinforce the growing trend in ultra-processed food consumption, also identified in other Brazilian populations^19,23^, potentially contributing substantially to the increased risk of non-communicable chronic diseases, including gastric cancer.

Statistical comparison showed that, for all patients, bean consumption was significantly higher than PCM and SUPF (p < 0.05), but lower than SSB (p < 0.05). Additionally, vegetable and fruit consumption was considerably higher compared to PCM and SUPF (p < 0.05). When comparing healthy foods (beans, legumes, vegetables, fruits) with unhealthy foods (PCM and SSB), in general, consumption of unhealthy foods was higher.

Gender analysis showed that women had slightly higher vegetable intake, while men consumed significantly more fruits compared to PCM and SUPF. However, men also had higher intake of PCM (30.23 g/day) and SSB (337.2 g/day) than women (Table 3). These differences may reflect distinct cultural or behavioral patterns, warranting further investigation. The associations observed align with studies linking excessive sugar intake to increased risk for various cancers, including gastric adenocarcinoma^28,29^.

Almahri et al.^29^ explored the relationship between dietary item consumption and the risk of developing gastric and pancreatic cancers, finding significant associations with high sugar intake, particularly sweets like candies and cookies, for gastric cancer. Intake of sugar-sweetened beverages is also associated with increased risk and mortality for several cancers, including gastric and colorectal, in a dose-dependent manner^30,31,32,33^. Consumption of these beverages increased overall cancer risk by 4% per 100 mL/day, while fruit juice consumption increased risk by 14% for the same increment^31,32^.

A prospective cohort study by Chazelas et al.^30^ inferred that consumption of sugar-sweetened beverages (sodas and fruit juices) was positively associated with overall cancer risk and mortality, with a subdistribution hazard ratio of 1.18 per 100 mL/day increase. High sugar intake may potentially increase cancer risk through mechanisms such as insulin-glucose dysregulation, oxidative stress, inflammation, and increased body fat^28^.

In the present study, the total mean intake of ultra-processed foods reached 540.2 g/day, with a median of 330.73 g/day, highlighting the prominence of this food group in the diet of patients with gastric adenocarcinoma. This dietary pattern is consistent with findings by Ferro et al.^7^, Isaksen & Dankel^34^, Santiago et al.^19^, and Silveira, Dos Santos, França^35^, who report excessive ultra-processed food consumption in the Brazilian population in general.

Conversely, healthy dietary patterns are associated with lower levels of inflammation, as fibers, omega-3 polyunsaturated fatty acids (PUFAs), vitamins C and E, fruits, and vegetables may exert protective effects. These dietary components contribute to reducing oxidative stress and controlling chronic inflammation, both processes intimately related to the pathophysiology of chronic diseases, including cancer^34,36,37^.

A critical point is the frequently observed inverse relationship between ultra-processed and sugary food consumption and intake of fresh foods such as fruits, vegetables, and legumes. Studies suggest that as ultra-processed food intake increases, the consumption of healthy foods tends to decrease proportionally^24^.

In this scenario, promoting dietary strategies that encourage regular consumption of fruits, vegetables, and greens becomes essential. Beyond providing essential micronutrients for metabolic homeostasis, these dietary choices may indirectly reduce ultra-processed food intake, contributing to chronic disease prevention and improving prognosis in individuals already diagnosed with neoplasms, such as gastric adenocarcinoma.

It was observed that dietary patterns can directly influence gene expression modulation and the tumor microenvironment in gastric adenocarcinoma. Differential gene expression analysis conducted on 16 patients, distributed between LEDRF and HEDRF, revealed approximately 48 differentially expressed genes. Six genes were prioritized for their potential biological role in gastric carcinogenesis, allowing a more in-depth investigation of their contribution to the distinct molecular signature observed between groups and possible modulation by specific dietary patterns.

Enrichment analysis and gene expression findings demonstrated that the main enriched pathways included diseases of metabolism and glycosylation, as well as alterations in keratan sulfate biosynthesis and degradation, surfactant metabolism, and water-soluble vitamin metabolism, potentially indicating tumor metabolic reprogramming that may modulate tumor progression and inflammatory response^38,39,40^.

In Group 0, overexpression of MAGEA3, IL6, and HCAR2 was observed, genes associated with cell proliferation, inflammation, and tumor microenvironment remodeling. In Group 1, underexpression of NLRP4, PGA3, and CTCFL was noted, genes involved in immune and metabolic homeostasis regulation. These findings reveal distinct molecular signatures between groups, suggesting that differences in dietary patterns may be related to modulation of biologically relevant pathways in gastric carcinogenesis.

Among differentially expressed genes, MAGEA3 and IL-6 exhibited marked overexpression in Group 0. This result reflects an apparently paradoxical biological behavior, since MAGEA3 is related to cell proliferation, immune evasion, and tumor progression^41^, while IL-6 is a classic pro-inflammatory cytokine involved in chronic inflammation and cancer progression, also considered a potential prognostic biomarker^42,43^. The strong activation of these genes in a lower dietary risk group may reflect the intrinsic complexity of GA, marked by high molecular heterogeneity.

In this context, the overexpression of MAGEA3 and IL-6 in the group with lower dietary risk exposure highlights the complexity of tumor biology and the possible influence of epigenetic and metabolic mechanisms independent of isolated dietary patterns. Moreover, MAGEA3 expression may be influenced by the genomic methylation profile, which can be modulated by dietary components such as polyphenols and folate^44,45,46^.

Additionally, pro-inflammatory diets high in processed meats are associated with higher IL-6 levels, while anti-inflammatory diets rich in vegetables and whole foods tend to reduce its expression. These findings reinforce the modulatory role of diet not only on metabolic and inflammatory processes but also on the expression of genes with significant clinical implications, suggesting potential targets for therapeutic and preventive strategies integrated with the dietary and molecular context of gastric adenocarcinoma.

In this study, HCAR2 exhibited overexpression in the LEDRF group, corresponding to lower exposure to ultra-processed foods. It is noteworthy that ligands of this receptor have epigenetic functions as histone deacetylase (HDAC) inhibitors, implicated in tumor progression^47^. Most evidence on HCAR2 relates to colorectal cancer. A recent study by Yu et al.^48^, using in vivo and in vitro models, demonstrated that selective agonists of this receptor, such as butyrate derived from fiber fermentation, induce immunomodulatory and anti-inflammatory responses, including increased CD8⁺ T cell anti-tumor activity and positive modulation of the tumor microenvironment with reduced inflammation.

Protective dietary patterns, such as those rich in vegetables, fruits, fiber, or ketogenic components, may increase the availability of niacin (vitamin B3) and ketone bodies, both natural ligands of HCAR2, which have been associated with inflammation modulation in the tumor microenvironment, contributing to a less permissive environment for cancer progression^47,48,49^.

In the group with higher exposure to dietary risk factors, significant underexpression of NLRP4, a gene known for its atypical regulatory role in inflammatory processes, was observed. NLRP4 acts as an inhibitor of inflammatory pathways, including type I interferon signaling via TBK1 degradation, and is involved in autophagy modulation^50^.

Underexpression of NLRP4 in this group may indicate weakened anti-inflammatory regulatory mechanisms in the tumor microenvironment, favoring the intensification of unbalanced innate immune responses. This dysregulation may contribute to chronic inflammation, which is known to be associated with gastric cancer progression^5,51^.

Still in the HEDRF group, PGA3 underexpression was observed, responsible for encoding a precursor of pepsin, a key digestive enzyme for protein hydrolysis. Its reduction aligns with evidence showing decreased PGA3 levels in atrophic gastritis and gastric adenocarcinoma^52^. Large-scale pan-cancer analyses using TCGA data^53^ also indicate that PGA3 shows reduced expression specifically in gastric adenocarcinoma^54^.

This expression profile is consistent with the inflammatory context characteristic of the HEDRF group, whose diet is high in salt, nitrites, processed meats, and other pro-inflammatory components, contributing to repeated gastric mucosa damage, favoring atrophy, intestinal metaplasia, and cellular instability, precursors of gastric carcinogenesis. Additionally, ultra-processed foods, rich in pro-oxidative and inflammatory agents, contribute to maintaining low-grade chronic inflammation^5,55^.

In the HEDRF group, CTCFL underexpression was also observed compared to the lower exposure group. Although hyporepressed, CTCFL may be actively transcribed in some cases, as under normal physiological conditions, its expression should be absent in somatic tissues^56,57^. CTCFL expression is documented in most cancerous tissues and cell lines. Normally restricted to specific cells and reactivated in cancer, it is considered a potential biomarker. Previous studies associate its presence with disease progression, advanced tumor stages, and worse prognosis in different cancers^58^.

These findings gain further relevance considering that diet can directly influence epigenetic mechanisms, including DNA methylation. Methyl-donor nutrients, such as folate, choline, methionine, and B vitamins, are essential for one-carbon metabolism, regulating S-adenosylmethionine availability, the main methyl donor for genomic methylation reactions. Their availability can modulate tumor suppressor or oncogene methylation, impacting expression and tumor behavior^59^.

Considering that vitamin, complex carbohydrate, and glycoconjugate metabolism is influenced by diet, it is plausible that patients dietary profiles, particularly higher exposure to ultra-processed foods and low nutrient intake, may contribute to modulating an inflammatory and metabolically adapted tumor microenvironment favoring neoplastic progression.

Clinical comparisons between HEDRF and LEDRF revealed no statistically significant differences in evaluated variables, including age, tumor staging, Lauren histological classification, and neoadjuvant treatment. Overall, findings suggest that, independent of overt clinical variables, distinct dietary exposures may be associated with modulation of biological mechanisms favoring an inflammatory and metabolically adapted tumor microenvironment in gastric adenocarcinoma. Lifestyle habits prior to diagnosis can significantly influence the occurrence of second primary cancers and long-term survival in patients with a history of gastric adenocarcinoma, reinforcing the importance of investigating modifiable factors such as dietary patterns^60^.

## Conclusion

This study demonstrated that GA, in the investigated population, presents an epidemiological profile characterized by higher occurrence in male individuals, advanced age, and unfavorable socioeconomic conditions, highlighting the influence of social determinants of health on the diagnosis and prognosis of the disease. The observed dietary patterns indicated insufficient consumption of unprocessed foods and a high intake of ultra-processed foods, particularly sugar-sweetened beverages and other ultra-processed foods, a characteristic that may contribute to a pro-oxidative and inflammatory tumor microenvironment.

Furthermore, the analysis of patients gene expression revealed approximately 48 differentially expressed genes between groups with lower versus higher exposure to dietary risk factors. The observed molecular signatures, together with pathway enrichment findings related to metabolism, glycosylation, and inflammatory response, suggest that diet may act as a modulator of epigenetic, metabolic, and immunological processes relevant to tumor progression. The results reinforce the need for integrated approaches in the prevention and management of gastric adenocarcinoma, ranging from dietary strategies and personalized nutritional interventions to the investigation of molecular biomarkers modulated by diet.

## Acknowledgment

The authors would like to thank the Oncology Research Center, the Human and Medical Genetics Laboratory, and the Anatomical Pathology Laboratory at João de Barros Barreto University Hospital (HUJBB – UFPA) for their invaluable technical and laboratory support. Our gratitude also goes to the High-Performance Computing Center (CCAD) at the Federal University of Pará for access to the Apollo 2000 cluster, which was crucial for our analyses. This work was supported by Fundação Amazonia de Amparo a Estudos e Pesquisas – FAPESPA (004/21), Conselho Nacional de Desenvolvimento Científico e Tecnológico – CNPq (313303/2021-5) and Ministério Público do Trabalho (11/12/2020 – Ids 372cfc4 and b7c1637).

## Data availability statement

The original contributions presented in the study are included in the article. Further inquiries can be directed to the corresponding author.

